# Inference of sexual selection from pairwise contests

**DOI:** 10.1101/2020.02.25.964973

**Authors:** Thomas Yu, Mehrnaz Afkhami, Andrew G. Clark

## Abstract

Sexual selection, whether mediated by male-male competition, female choice, male choice, or other factors influencing reproductive outcomes, can be usefully framed as a series of pairwise contests. Experimental evolutionary biologists have long used male-choice and female-choice experiments to quantify genotype-specific mating success, a key component of fitness. Here, we consider *Drosophila* mate-choice experiments and apply the Bradley-Terry model, which has been explicitly designed to quantify the merits of contestants and establish a ranking based on results from pairwise contests. The Bradley–Terry model can also be used to predict the probabilities of outcomes for contests that have not yet taken place. Leave-one-out validation allows us to assess both model fit and predictive performance for unobserved contests. After applying these methods to *Drosophila* mate-choice experiments, we interpret the results in terms of sexual selection and discuss the implications of deviations from a well-fitting model.

## Introduction

Since Darwin (1859, 1896) first introduced and formalized the idea of sexual selection, it has come to be seen as a potent driver of evolutionary change (Andersson 1994). The genetic variance for mating success appears in many organisms to exceed the genetic variance for other classical components of fitness, such as viability (Bundgaard and Christiansen 1972). This observation establishes a key role for sexual selection in evolution and motivates the need for sound empirical methods for quantifying it. A wide variety of empirical approaches have been applied to quantify sexual selection. One of the most common experimental protocols is to present virgin individuals of one sex with a pair of individuals of the opposite sex and observe the first mating that occurs (Arbuthnot et al. 2017; Dechaume-Moncharmont et al. 2013; Herdman et al. 2004). Although such pairwise contests are somewhat artificial and may not fully reflect natural conditions, many trials in aggregate can yield robust inferences of genotype-specific mating success.

The Bradley–Terry model (Bradley and Terry 1952) provides a powerful statistical framework for analyzing outcomes of pairwise mating contests by estimating an underlying “ability” parameter for each individual or genetic line. These parameters are inferred from the collection of observed contests, allowing the model to integrate information across all pairwise comparisons and to generate probabilistic predictions for contests that have not been observed (Cattelan 2012; Király and Qian 2017; Fang et al. 2026). The method remains effective even when not all possible pairwise matches have been performed. This approach is particularly well suited to laboratory mate-choice experiments, where replicated pairwise trials can help stabilize inference from inherently noisy data on mating success or dominance. The Bradley–Terry model specifies a generative process for expected mating frequencies, yielding coherent estimates of relative mating abilities and enabling formal tests of whether outcomes are consistent with a simple hierarchical ranking. Moreover, the framework facilitates quantitative assessment of experimental design, such as the number of trials required for reliable inference, and improves power to detect differences among genotypes. When mating success is the primary fitness component, the resulting ability estimates can be interpreted as relative fitness measures. The Bradley–Terry model has been especially useful in establishing associations between mating success and phenotypic and behavioral attributes. Mating success was positively associated with morphological and signaling traits in several lizard species (Abalos et al. 2016, 2024; Gilbert et al. 2020; Stuart-Fox et al. 2006). On the other hand, traits assumed to confer mating advantages, such as bite force, sprint speed and endurance, were convincingly shown with Bradley-Terry results to have no association with mating success in skinks (Noble et al. 2021). Haley et al. (1994) applied the Bradley-Terry model to establish the strong relation between body size and mating success in elephant seal males, and Jaworski et al. (2018) showed that larger females were preferred mates in the red-backed salamander. In all these examples, mating success or social dominance was associated with phenotypes, whereas stronger inference about sexual selection requires demonstrating that mating success depends on heritable genetic differences among individuals or lines.

In this study, we fit the Bradley–Terry model to mate-choice data from Arbuthnott et al. (2017) and examine differences in ability estimates across genotypes and evaluate overall model fit. Rather than associate mating success with phenotypic attributes, we quantified differences in mating success among genetic lines, thereby demonstrating line-specific, heritable variation in mating success under these experimental conditions. The good fit observed here supports the interpretation that, in this experimental setting, sexual selection can be summarized by a ranking of the genotypes tested, such that outcomes of pairwise contests can be predicted from their estimated abilities.

## Materials and Methods

### Experimental design

To apply the Bradley-Terry model to a mate-choice experiment, we needed a study that considered many pairwise competitions among a collection of multiple lines. *Drosophila* mate-choice data were generously provided by Devin Arbuthnott from Arbuthnott et al. (2017). That study was designed as a male-choice test to determine whether male choice among pairs of female genotypes exhibits transitivity. Note that although trials with a single male and two females are often called male-choice tests, the difference in estimated ‘ability’ may be driven by any combination of male choice, female attractiveness, and female willingness to mate. Traditionally, female Drosophila have been viewed as either accepting or rejecting male advances, and most mate-choice experiments in this genus have therefore been conducted as female-choice tests (Andersson 1994). Part of the motivation of Arbuthnott et al. (2017) was to assess whether male choice is an important phenomenon in flies. We used the data from pairwise contests with Oregon-R males and females from eleven lines from the *Drosophila* Genetic Reference Panel (DGRP): 138, 313, 362, 399, 517, 555, 707, 714, 738, 774, and 786.

Because each line is a distinct inbred genotype, we use the terms “line” and “genotype” interchangeably. Here we briefly describe how the data were collected: in each trial, two females, one from each of a pair of DGRP lines, were placed together in a single vial with a male from an Oregon-R stock. In each trial, the two females were dusted with differently colored fluorescent powder to allow them to be distinguished. Matings were randomized across time.

Altogether, there were 55 distinct pairwise tests, and a total of 550 mating trials were conducted. The line number of the female that successfully mated within a 2-hour observational period was recorded. Among the 550 mating trials, 392 concluded with a successful mating.

### Statistical model

The basic modeling challenge is to estimate, a vector of “ability” scores, λ*_i_*, for *i* = 1 to 11, based on the results of a series of pairwise contests among our 11 genetic lines of flies. With these λ*_i_* estimates, the formula for the probability of line *i* being chosen for mating over line *j* is

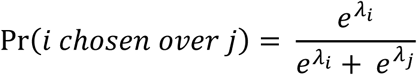

These λ*_i_* mating ability scores could, in principle, serve as a metric of relative fitness under sexual selection. Given a matrix of observed wins and losses for each pairwise contest, Bradley and Terry (1952) derived maximum likelihood estimators for the *λ_i_*.

### Parameter estimation

We used R version 3.5.2 for the data analysis (R Core Team 2018). The R package readxl (Wickham and Bryan 2019) was used to import the data, and the BradleyTerry2 package (Turner and Firth 2012) was used to fit the Bradley–Terry model and estimate mating abilities and their standard error. We used ggplot2 to visualize the data (Wickham 2016).

### Model validation

#### Deviance statistics

The R package reports both null deviance and residual deviance to assess model fit. The residual deviance quantifies goodness-of-fit by measuring the amount of variation remaining unexplained after the model has been fitted. A smaller deviance value indicates a better fitting model. The null deviance corresponds to a model with no fitted parameters, whereas the residual deviance corresponds to a model including all line-specific parameters. The difference between the residual and null deviances yields a test statistic that approximately follows χ^2^ distribution with 10 degrees of freedom (11 lines minus one constraint). We use a chi-square test to compute *P*-values and assess whether the fitted model provides a significantly better fit than the null model.

#### Leave-one-out tests

We assessed model performance by systematically excluding results for each pairwise contest between two lines. We fit the Bradley–Terry model to the reduced dataset and used it to predict the outcome probability for the excluded pair. We then compared predicted mating counts with observed mating counts using a chi-square test (with one degree of freedom) to evaluate predictive accuracy. With 11 lines there are 55 pairs, yielding 55 chi-square tests, each with one degree of freedom.

#### Transitivity tests

One key assumption of the Bradley-Terry model is that the contests exhibit transitivity. Transitivity, an indicator of “rational” behavior (Arbuthnott et al. 2017), occurs when, if A beats B, and B beats C, then A beats C. Weak stochastic transitivity occurs when, given Pr(A beats B) ≥ 0.5 and Pr(B beats C) ≥ 0.5, then Pr(A beats C) ≥ 0.5 (Houston 1991). Strong stochastic transitivity occurs when, given Pr(A beats B) ≥ 0.5 and Pr(B beats C) ≥ 0.5, then Pr (A beats C) ≥ max[Pr(A beats B), Pr(B beats C)] (Houston 1991). Equations 2a and 2b hold true only under strong stochastic transitivity. A good fit to the Bradley–Terry model implies not only transitivity but also accurate quantitative prediction of mating probabilities. Transitivity alone, without a parametric model like Bradley–Terry, does not provide quantitative predictions of mate-choice outcomes.

Let A, B, and C denote three lines. We enumerate all trios and order them such that A > B and B > C, where “>” indicates a greater number of matings. After temporarily excluding the A vs. C result, we fit a Bradley–Terry model and predict the probability that a male chooses a female from line A over line C. We compare this prediction to the observed proportion of trials in which A is chosen over C. We conduct a binomial test to generate a *P*-value for each of the 165 tests (the number of distinct trios of lines possible from a set of 11 lines).

For each leave-one-out test we get observed and expected counts of the two genotypes of females that mated and calculate the binomial probability of the difference. We compare the observed *P*-values from the binomial tests to the expected uniform distribution. This comparison reveals deviations from the expected null distribution.

## Results

The Bradley–Terry model revealed clear differences in mating ability among the 11 DGRP female genotypes tested in pairwise contests with Oregon-R males. Estimated abilities were stable with increasing numbers of contests and showed good overall agreement between observed and predicted outcomes at both the pair and trio levels.

### Bradley–Terry abilities and genotype ranking

Fitting the Bradley–Terry model to the 55 pairwise contests yielded distinct ability estimates for each line, indicating substantial heterogeneity in mating success (Figure 1 and Table 1). Line 738 was arbitrarily chosen as the baseline, so abilities for the other lines are expressed relative to line 738. Lines 517 and 313 had the lowest estimated abilities, whereas lines 774, 738 (baseline), and especially 362 had much higher abilities. For example, the model predicts that 362 will be chosen over 738 with probability *e*^0.905^/(*e*^0.905^+*e*^0^) = 0.712. By contrast, a pair of mid-ranking lines with similar mating abilities, such as 555 and 714, yields a more even match, with the predicted probability *e*^−1.542^/(*e*^−1.542^+*e*^−1.1229^) = 0.420. Thus, the fitted abilities provide a quantitative ranking of genotypes that can be interpreted directly in terms of expected mating success in any pairwise contest under these experimental conditions.

**Figure 1.**
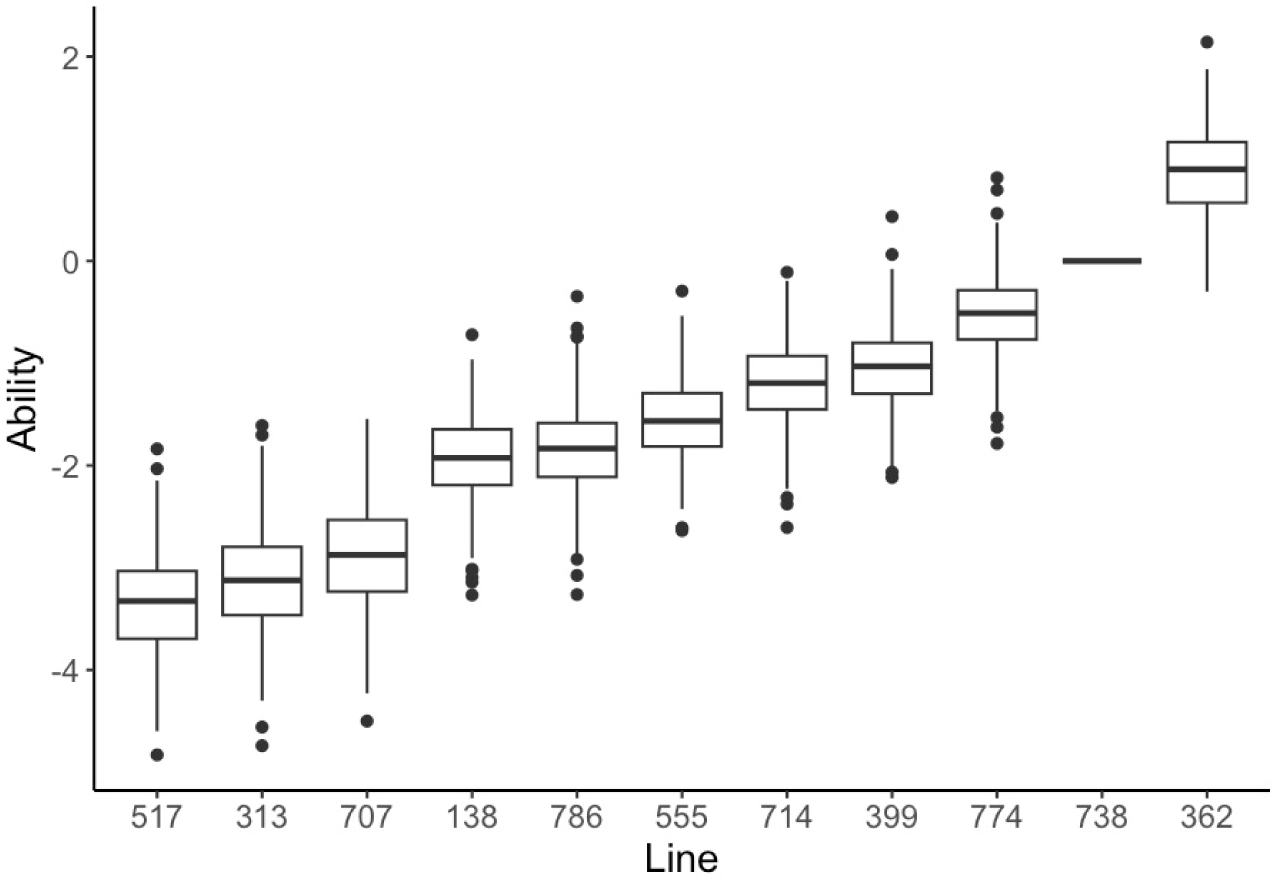
Bradley-Terry mating abilities (λ*_i_*) estimated for each line using the Bradley-Terry model. Data were from 550 pairwise contests of all possible pairs of females challenged with a single Oregon-R male.

**Table 1.** Female mating ability estimates (λ*_i_* and standard errors) of the 10 *Drosophila* DGRP lines (normalized to line 738) according to the Bradley-Terry model. From these mating ability estimates, the prediction that line *i* will be favored over line *j* in this pairwise contest is 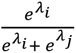.

| Line | Bradley-Terry Ability |
| --- | --- |
| 517 | -3.365 (0.54) |
| 313 | -3.141 (0.48) |
| 707 | -2.852 (0.47) |
| 138 | -1.918 (0.41) |
| 786 | -1.836 (0.42) |
| 555 | -1.542 (0.39) |
| 714 | -1.229 (0.39) |
| 399 | -1.037 (0.37) |
| 774 | -0.521 (0.38) |
| 738 | 0.00 |
| 362 | 0.905 (0.42) |

### Stability of ability estimates and rankings

To assess how much data are needed for reliable inference, we added contests sequentially in random order and recomputed the Bradley–Terry abilities after each additional set of mating trials (Figure 2). Ability estimates for all lines changed rapidly at small sample sizes but stabilized after roughly 200 contests, with only minor fluctuations as additional data were added. In contrast, the rank order of lines sometimes changed even late in the sequence when estimated abilities were very similar. This indicates that robust parameter estimation is achievable with modest sample sizes, but fine-scale ranking among nearly equal genotypes remains sensitive to sampling noise.

**Figure 2.**
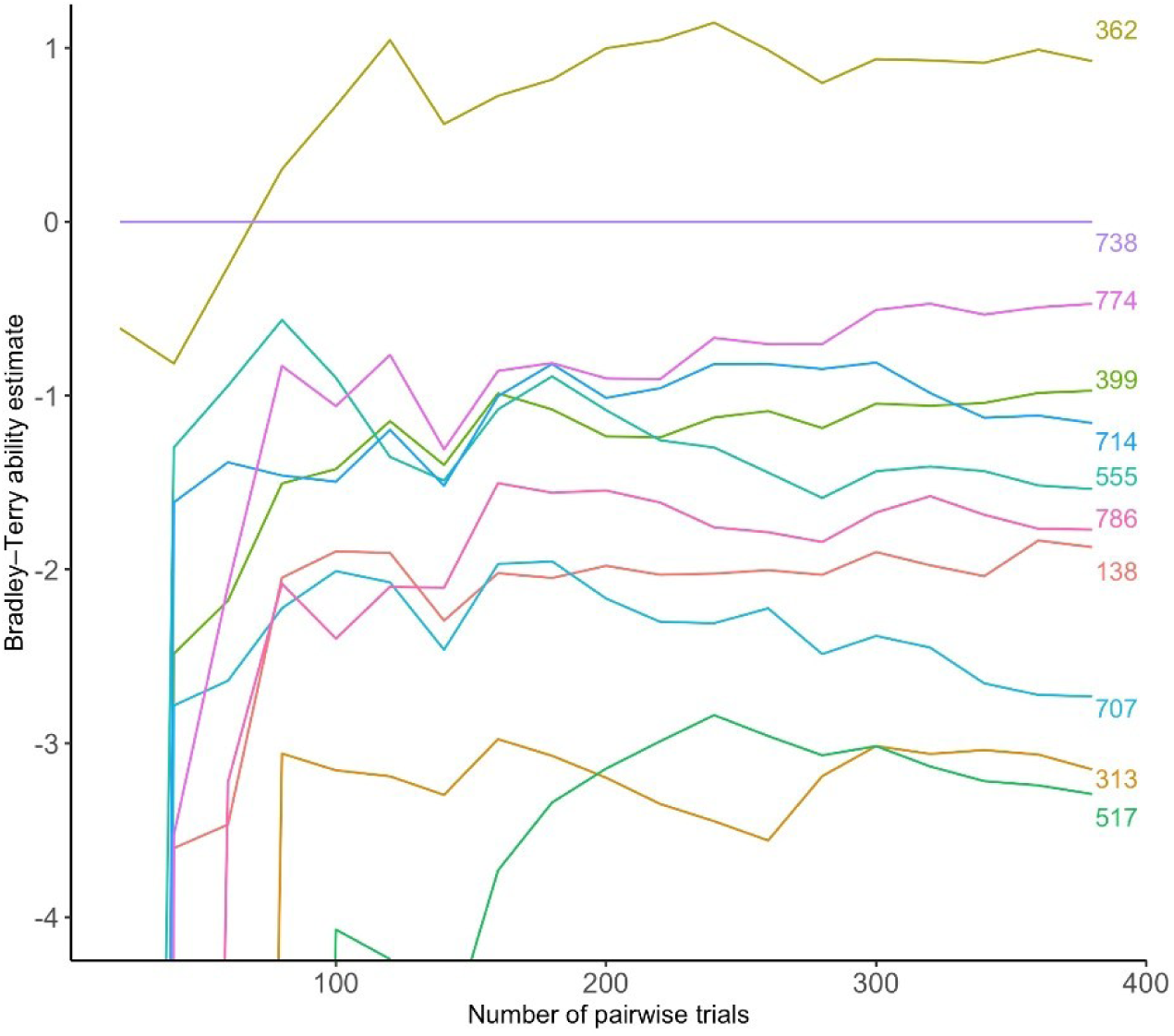
Bradley-Terry abilities estimated from subsets of the data, showing how the estimates stabilize as the number of trials grows. Pairwise contests were chosen at random (without replacement), and Bradley-Terry mating abilities were recalculated after each additional set of trials. Abilities are normalized to line 738, yielding a horizontal line at 0 for this line. Shown is a typical replicate of the resampling process. In this replicate, after 220 pairwise tests, the rank order of the ability estimates remained relatively stable.

### Overall goodness-of-fit

Deviance-based diagnostics indicated a strong improvement of the Bradley–Terry model over a null model with equal mating abilities. The null deviance was 249.07 with 55 degrees of freedom, whereas the residual deviance of the fitted model was 56.70 with 45 degrees of freedom, corresponding to an extremely small chi-square *P*-value (6.29 × 10^−36^). This large reduction in deviance shows that models allowing line-specific mating abilities provide a substantially better explanation of the observed contest outcomes than a model with no differences among genotypes.

### Leave-one-out prediction of pairwise contests

We next evaluated predictive performance at the level of individual pairs by leave-one-out cross-validation. For each of the 55 line pairs, we removed all contests between that pair, refit the model to the remaining data, and used the refitted model to predict the probability that one line would be chosen over the other. Comparing predicted probabilities to observed win proportions with binomial tests showed that most pairs lay close to the one-to-one line (Figure 3), and only one of the 55 tests yielded a significant deviation at α = 0.05. This pattern is consistent with the number of significant tests expected by chance under a well-fitting model and indicates that the Bradley–Terry parameters have meaningful predictive power for contests not used in fitting.

**Figure 3.**
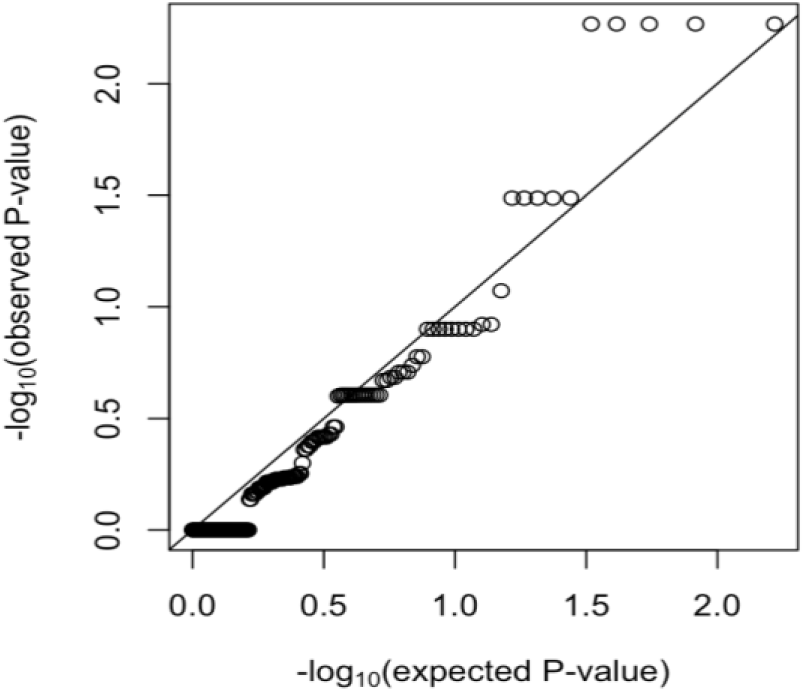
Results of leave-one-out tests of the ability of the Bradley-Terry model to predict mate choices when blinded to the data of one of the 55 pairwise tests. See text for details.

### Stochastic transitivity and trio-based tests

To examine transitivity, we considered all ordered trios of lines A, B, and C such that A beat B and B beat C in the observed data and then temporarily excluded the A–C contests. For each of the 165 such trios, we used the fitted Bradley–Terry model to predict the probability that A would be chosen over C and compared this prediction to the observed proportion using a binomial test. Ten of the 165 tests were significant at α = 0.05, close to the fraction expected by chance. These results indicate that the mate-choice data are broadly consistent with the stochastic transitivity assumptions of the Bradley–Terry model and show no clear pattern of systematic violations.

### Integrated assessment of model performance

Across all diagnostics, the Bradley–Terry model provides a coherent and well-supported description of these *Drosophila* mate-choice data. Ability estimates yielded a stable ranking of genotypes and explain most of the deviance relative to a null model, and leave-one-out tests at both the pair and trio levels show that predicted mating probabilities closely match observed outcomes. Together, these findings justify interpreting the estimated mating abilities as robust quantitative measures of mating success for these inbred lines in this experimental setting and motivate their use as proxies for sexual fitness components in subsequent theoretical analyses.

## Discussion

Evolutionary biologists have often used mate choice tests to assess differences among genotypes in mating ability, an essential component of fitness. For the most part, inference about sexual selection from aggregated pairwise tests has relied on simple heuristics or ad hoc statistics rather than on explicit generative models. Here, we have shown that the Bradley–Terry framework provides a rigorous way to estimate genotype-specific scores that serve as reasonable proxies for relative mating success. These models make specific assumptions about the data and the situation to which they are applied (Shev et al. 2014). First, the model assumes that there is a true underlying ranking of participants and that contest outcomes are transitive. Contests are assumed to be independent of each other, and outcomes are assumed not to depend on broader context or past contests. Given the adequate fit of these Drosophila mate-choice data to the Bradley–Terry model, these assumptions appear broadly consistent with the observed outcomes, at least in this constrained laboratory environment.

There are several advantages to the application of the Bradley-Terry model to pairwise contest data in the context of evolutionary biology. A formal parametric model allows the assessment of the overall goodness-of-fit of the model to the data. This allows the identification of lines or assays that deviate from the model. Additionally, these analyses inform data quality, particularly whether the experiment is sufficiently large and consistently executed to yield stable and robust parameter estimates (Figure 2). Finally, the parameter estimates from the model explicitly present the researcher with the ability to predict the outcomes of other pairwise tests, allowing a powerful form of cross-validation.

This study also raises interesting questions about the old “unit of selection” debate (Lewontin 1970). Sexual selection can be modeled with a fitness parameter for each genotype, so that the outcome of a particular mating trial depends only on the parameters of the two genotypes involved (e.g., Equation 1). In this case, the unit of selection is the single genotype, and the model has as many parameters as there are genotypes. The good fit to the Bradley-Terry model implies that these mate-choice test results are consistent with this simple genotype-based model of sexual selection. If the model had not fit well, one plausible interpretation would be that there are specific interactions between particular pairs of females, in which case the relevant unit of selection might shift from single genotypes to pairs of genotypes. Outcomes of mating trials could then be modeled with a matrix of parameters whose entries represent all possible genotype pairs, extending the Bradley–Terry framework to include pair-specific effects.

Our analysis implicitly assumes that the Bradley–Terry parameters are invariant across time and other contexts. Violations of these assumptions are particularly problematic if one aims to use fitted Bradley–Terry parameters to predict future allele frequency dynamics. First, the parameter estimates are likely to depend on environmental conditions, including diet, temperature, and degree of crowding. A fly’s mating performance in the wild is not determined by a single virgin choice test, but instead reflects outcomes across many trials, allowing for experience-dependent changes in behavior and choice. Finally, the fly lines used in the experiment are highly inbred, so after the first generation of matings many classes of heterozygous offspring would be produced, complicating any direct mapping from line-specific abilities to allele frequency trajectories. Thus, a key limitation of this experiment is that the estimated mating abilities cannot be directly used to predict allele dynamics in an outbred population.

The Bradley-Terry model, and related approaches that fit large numbers of pairwise contests (Hamilton et al. 2023; Fang et al. 2026), have multiple fruitful applications in evolutionary biology. Joint analysis of male-choice and female-choice experiments would allow assessment of the relative contributions of male behavior, female behavior, and ‘luck’ to mating outcomes (Gilbert and Wells 2019). At present, female choice is generally regarded as the dominant factor in *Drosophila* sexual selection, but recent studies have shown that male choice is repeatable and biologically meaningful (Arbuthnott et al. 2017; Edward and Chapman 2012). Finally, we note that inference and prediction in the context of pairwise tests is an area of active development, including Bayesian methods (Leonard 1977; Issa Mattos and Martins Silva Ramos 2022; Schaumberger and Tutz 2019; Gilbert and Wells 2019) and machine learning approaches (Menke and Martinez 2008). Extensions that accommodate multiple covariates have been developed (Casalicchio et al. 2015; Hatzinger and Dittrich 2012). These methods can even accommodate multi-dimensional attributes of the competing individuals, making them even more useful in a broad collection of evolutionary applications.

## Authors’ contributions

AGC conceived the study, MA and TY performed the coding to format the data and obtain parameter estimates, and all authors wrote and edited the paper.

## Acknowledgements

This work was supported by NIH R01 HD059060 to AGC and M. F. Wolfner. We thank Martin Wells for insightful comments as we developed the ideas and approaches used in this manuscript. Ana Marija Jakšić, Arvid Ågren, Ian Caldas, Jullien Flynn, Manisha Munasinghe and Ulises Rosas Puchuri provided helpful suggestions for the manuscript.

## Data accessibility

All scripts developed in this paper will be available on GitHub upon publication.

## SUPPLEMENTARY TABLES

**Table S1.** Matrix of expected probabilities from the Bradley-Terry model. *P_ij_* is the probability that line *i* will be selected over line *j*. Probability ± 1 standard error. *z*-test for H0: *P_ij_* = 0.5(ns: P > 0.05, *: P < 0.05, **: P < 0.01, ***: P < 0.001) Significance codes appear in parentheses.

|  | 738 | 399 | 313 | 555 | 138 | 774 | 517 | 786 | 714 | 362 | 707 |
| --- | --- | --- | --- | --- | --- | --- | --- | --- | --- | --- | --- |
| 738 | 1.00 | 0.74±<br>0.07<br>(***) | 0.96±<br>0.02<br>(***) | 0.82±<br>0.06<br>(***) | 0.87±<br>0.05<br>(***) | 0.63±<br>0.09<br>(ns) | 0.97±<br>0.02<br>(***) | 0.86±<br>0.05<br>(***) | 0.77±<br>0.07<br>(***) | 0.29±<br>0.09<br>(*) | 0.95±<br>0.02<br>(***) |
| 399 | 0.26±<br>0.07<br>(***) | 1.00 | 0.89±<br>0.06<br>(***) | 0.62±<br>0.13<br>(ns) | 0.71±<br>0.11<br>(ns) | 0.37±<br>0.12<br>(ns) | 0.91±<br>0.05<br>(***) | 0.69±<br>0.12<br>(ns) | 0.55±<br>0.13<br>(ns) | 0.13±<br>0.06<br>(***) | 0.86±<br>0.07<br>(***) |
| 313 | 0.04±<br>0.02<br>(***) | 0.11±<br>0.06<br>(***) | 1.00 | 0.17±<br>0.09<br>(***) | 0.23±<br>0.11<br>(*) | 0.07±<br>0.04<br>(***) | 0.56±<br>0.18<br>(ns) | 0.21±<br>0.11<br>(**) | 0.13±<br>0.07<br>(***) | 0.02±<br>0.01<br>(***) | 0.43±<br>0.17<br>(ns) |
| 555 | 0.18±<br>0.06<br>(***) | 0.38±<br>0.13<br>(ns) | 0.83±<br>0.09<br>(***) | 1.00 | 0.59±<br>0.14<br>(ns) | 0.26±<br>0.11<br>(*) | 0.86±<br>0.08<br>(***) | 0.57±<br>0.14<br>(ns) | 0.42±<br>0.14<br>(ns) | 0.08±<br>0.04<br>(***) | 0.79±<br>0.1<br>(**) |
| 138 | 0.13±<br>0.05<br>(***) | 0.29±<br>0.11<br>(ns) | 0.77±<br>0.11<br>(*) | 0.41±<br>0.14<br>(ns) | 1.00 | 0.2±0<br>.09<br>(***) | 0.81±<br>0.11<br>(**) | 0.48±<br>0.15<br>(ns) | 0.33±<br>0.13<br>(ns) | 0.06±<br>0.03<br>(***) | 0.72±<br>0.13<br>(ns) |
| 774 | 0.37±<br>0.09<br>(ns) | 0.63±<br>0.12<br>(ns) | 0.93±<br>0.04<br>(***) | 0.74±<br>0.11<br>(*) | 0.8±0<br>.09<br>(***) | 1.00 | 0.95±<br>0.03<br>(***) | 0.79±<br>0.1<br>(**) | 0.67±<br>0.12<br>(ns) | 0.19±<br>0.09<br>(***) | 0.91±<br>0.05<br>(***) |
| 517 | 0.03±<br>0.02<br>(***) | 0.09±<br>0.05<br>(***) | 0.44±<br>0.18<br>(ns) | 0.14±<br>0.08<br>(***) | 0.19±<br>0.11<br>(**) | 0.05±<br>0.03<br>(***) | 1.00 | 0.18±<br>0.1<br>(**) | 0.11±<br>0.06<br>(***) | 0.01±<br>0.01<br>(***) | 0.37±<br>0.17<br>(ns) |
| 786 | 0.14±<br>0.05<br>(***) | 0.31±<br>0.12<br>(ns) | 0.79±<br>0.11<br>(**) | 0.43±<br>0.14<br>(ns) | 0.52±<br>0.15<br>(ns) | 0.21±<br>0.1<br>(**) | 0.82±<br>0.1<br>(**) | 1.00 | 0.35±<br>0.13<br>(ns) | 0.06±<br>0.03<br>(***) | 0.73±<br>0.12<br>(ns) |
| 714 | 0.23±<br>0.07<br>(***) | 0.45±<br>0.13<br>(ns) | 0.87±<br>0.07<br>(***) | 0.58±<br>0.14<br>(ns) | 0.67±<br>0.13<br>(ns) | 0.33±<br>0.12<br>(ns) | 0.89±<br>0.06<br>(***) | 0.65±<br>0.13<br>(ns) | 1.00 | 0.11±<br>0.06<br>(***) | 0.84±<br>0.09<br>(***) |
| 362 | 0.71±<br>0.09<br>(*) | 0.87±<br>0.06<br>(***) | 0.98±<br>0.01<br>(***) | 0.92±<br>0.04<br>(***) | 0.94±<br>0.03<br>(***) | 0.81±<br>0.09<br>(***) | 0.99±<br>0.01<br>(***) | 0.94±<br>0.03<br>(***) | 0.89±<br>0.06<br>(***) | 1.00 | 0.98±<br>0.01<br>(***) |

**Table S2.** Matrix of mating success percentages from the data, along with tests of significance of goodness-of-fit of the leave-one-out tests. *D_ij_* is the observed percentage of times line *i* was selected over line *j* in the pairwise mating trails, and *P_ij_* is the model prediction for the same quantity. Binomial test for H0: *D_ij_* =*P_ij_* (ns: p>0.05, ∗: p<0.05, ∗∗: p<0.01, ∗∗∗: p <0.001). Significance codes appear in parentheses.

|  | 738 | 399 | 313 | 555 | 138 | 774 | 517 | 786 | 714 | 362 | 707 |
| --- | --- | --- | --- | --- | --- | --- | --- | --- | --- | --- | --- |
| 738 | NA | 0.9<br>(ns) | 0.88<br>(ns) | 0.78<br>(ns) | 0.88<br>(ns) | 0.56<br>(ns) | 1 (ns) | 0.83<br>(ns) | 0.71<br>(ns) | 0.4<br>(ns) | 0.83<br>(ns) |
| 399 | 0.1<br>(ns) | NA | 0.88<br>(ns) | 0.56<br>(ns) | 0.75<br>(ns) | 0.33<br>(ns) | 1 (ns) | 0.89<br>(ns) | 0.67<br>(ns) | 0 (ns) | 0.83<br>(ns) |
| 313 | 0.13<br>(ns) | 0.13<br>(ns) | NA | 0.14<br>(ns) | 0.25<br>(ns) | 0 (ns) | 0 (ns) | 0 (ns) | 0.14<br>(ns) | 0.1<br>(ns) | 0.67<br>(ns) |
| 555 | 0.22<br>(ns) | 0.44<br>(ns) | 0.86<br>(ns) | NA | 0.29<br>(ns) | 0.38<br>(ns) | 1 (ns) | 0.5<br>(ns) | 0.43<br>(ns) | 0 (ns) | 0.86<br>(ns) |
| 138 | 0.13<br>(ns) | 0.25<br>(ns) | 0.75<br>(ns) | 0.71<br>(ns) | NA | 0 (ns) | 1 (ns) | 0.33<br>(ns) | 0.17<br>(ns) | 0 (ns) | 0.86<br>(ns) |
| 774 | 0.44<br>(ns) | 0.67<br>(ns) | 1 (ns) | 0.63<br>(ns) | 1 (ns) | NA | 0.5<br>(*) | 0.83<br>(ns) | 0.43<br>(ns) | 0.22<br>(ns) | 1 (ns) |
| 517 | 0 (ns) | 0 (ns) | 1 (ns) | 0 (ns) | 0 (ns) | 0.5<br>(*) | NA | 0.4<br>(ns) | 0.2<br>(ns) | 0 (ns) | 0 (ns) |
| 786 | 0.17<br>(ns) | 0.11<br>(ns) | 1 (ns) | 0.5<br>(ns) | 0.67<br>(ns) | 0.17<br>(ns) | 0.6<br>(ns) | NA | 0.43<br>(ns) | 0 (ns) | 1 (ns) |
| 714 | 0.29<br>(ns) | 0.33<br>(ns) | 0.86<br>(ns) | 0.57<br>(ns) | 0.83<br>(ns) | 0.57<br>(ns) | 0.8<br>(ns) | 0.57<br>(ns) | NA | 0.2<br>(ns) | 0.57<br>(ns) |
| 362 | 0.6<br>(ns) | 1 (ns) | 0.9<br>(ns) | 1 (ns) | 1 (ns) | 0.78<br>(ns) | 1 (ns) | 1 (ns) | 0.8<br>(ns) | NA | 1 (ns) |
| 707 | 0.17<br>(ns) | 0.17<br>(ns) | 0.33<br>(ns) | 0.14<br>(ns) | 0.14<br>(ns) | 0 (ns) | 1 (ns) | 0 (ns) | 0.43<br>(ns) | 0 (ns) | NA |

**Figure S1.**
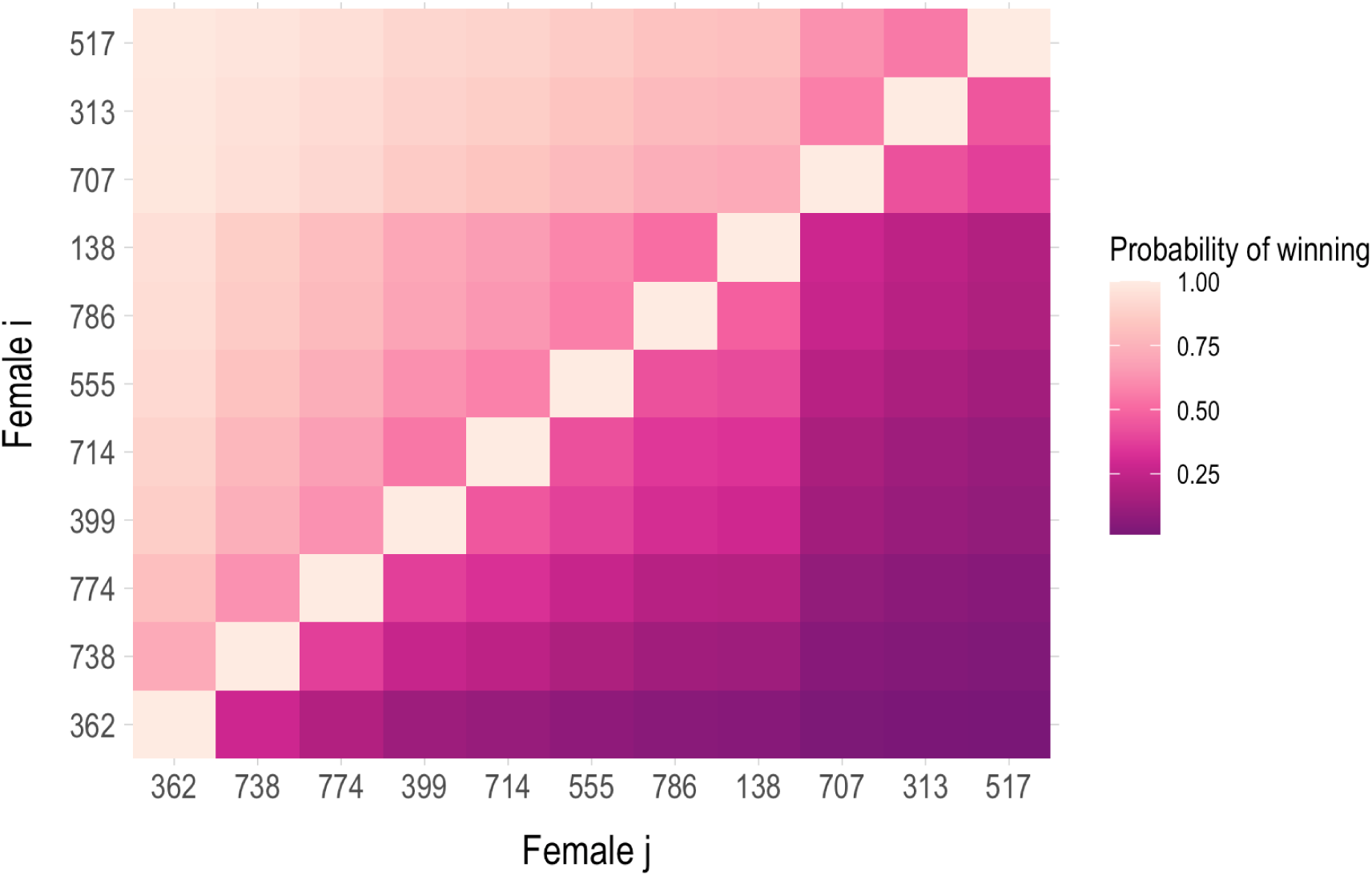
Heatmap showing, for all 55 pairwise mate-choice tests, the estimated probability that female from line *i* is chosen over the female from line *j* (upper triangle) and the lower triangle shows the probability that the female from line *j* is chosen over the female from line *i*.

